# Protein interaction network analysis reveals growth conditions-specific crosstalk between chromosomal DNA replication and other cellular processes in *E. coli*

**DOI:** 10.1101/2021.12.08.471875

**Authors:** Joanna Morcinek-Orłowska, Beata Maria Walter, Raphaël Forquet, Dominik Cysewski, Maxime Carlier, Sam Meyer, Monika Glinkowska

## Abstract

*E. coli* and many other bacterial species can alter their cell cycle according to nutrient availability. Under optimal conditions bacteria grow and divide very fast but they slow down the cell cycle when conditions deteriorate. This adaptability is underlined by mechanisms coordinating cell growth with duplication of genetic material and cell division. Several mechanisms regulating DNA replication process in *E. coli* have been described with biochemical details so far. Nevertheless we still don’t fully understand the source of remarkable precision that allows bacterial cells to coordinate their growth and chromosome replication. To shed light on regulation of *E. coli* DNA replication at systemic level, we used affinity purification coupled with mass spectrometry (AP-MS) to characterize protein-protein interactions (PPIs) formed by key *E. coli* replication proteins, under disparate bacterial growth conditions and phases. We present the resulting dynamic replication protein interaction network (PIN) and highlight links between DNA replication and several cellular processes, like outer membrane synthesis, RNA degradation and modification or starvation response.

**Importance:** DNA replication is a vital process, ensuring propagation of genetic material to progeny cells. Despite decades of studies we still don’t fully understand how bacteria coordinate chromosomal DNA duplication with cell growth and cell division under optimal and stressful conditions. At molecular level, regulation of processes, including DNA replication, is often executed through direct protein-protein interactions (PPIs). In this work we present PPIs formed by the key *E. coli* replication proteins under three different bacterial growth conditions. We show novel PPIs with confirmed impact on chromosomal DNA replication. Our results provide also alternative explanations of genetic interactions uncovered before by others for *E*.*coli* replication machinery.

## Introduction

At molecular level, most of biological functions are performed by proteins. Composite cellular activities are usually carried out by various sets of proteins organized into protein complexes, rather than by proteins acting alone. Formation of a complex, which is mediated by protein-protein interactions between their components, ensures coordination of processes and channeling of substrates. Often, protein assemblies gain additional features in comparison to their individual units, so that ultimately a protein complex means more than a simple sum of its parts. The entirety of protein-protein interactions in a cell constitutes a protein interaction network (1, 2). Recent studies on protein network properties revealed their general characteristics like connectivity, modularity and dynamics (3). It was shown that most proteins (nodes) within a network form limited number of connections (edges), but relatively small number of proteins form highly connected nodes called hubs. Moreover, proteins responsible for certain autonomous cellular activities are organized into well connected modules, typically forming few interactions between each other (1). Those intramodular connections may play a pivotal role in coordinating various cellular functions or in concerted response of cells to external stimuli. However, protein-protein interactions are also differentiated with respect to affinity of interactants and assembly on/off rate, ranging from very stable to transient ones. Hence, protein complexes are usually understood as stable multimolecular machines that act at the same time and place, whereas modules contain proteins that participate in a particular process, but their interactions are spatiotemporally regulated (4–6). Dynamics of proteins and PPIs underlies transformations of PINs under different conditions and reflects cellular responses to environmental cues, cell cycle or developmental stages. When conditions change, certain functional modules may continue, change composition, dissipate or fuse, forming additional connections (7–10). Uncovering dynamics of such condition-specific protein sub-networks dedicated to a certain process can facilitate understanding of their regulation at systemic level.

Bacterial cell cycle is a sequence of events directed by spatiotemporally regulated protein complexes. It consists of concurrent and interrelated processes: cell growth, chromosomal DNA replication and segregation culminated with cell division (11). For fast-growing bacteria like *E. coli*, the time needed for synthesis of full chromosomal copy exceeds the interval between subsequent divisions and, to cope with that, all cell cycle stages occur simultaneously (Fig. 1A)(12). Thus, bacterial cell may contain several replicating chromosomes at different replication stages, however still one initiation of replication per cell cycle rule is held (12). Coordination of DNA replication with cell growth and cell division in bacteria is still insufficiently understood. It is widely accepted that DNA replication starts at a certain cell volume/chromosomal origin ratio and that all origins of replication (*oriC*) present in the cell fire simultaneously (Fig. 1B)(13–15), but molecular mechanism behind this dependence remains uncertain. The key components of the mentioned control principles are certainly the DnaA protein that initiates a sequence of events at *oriC* leading to replication complex formation, as well as its regulators DiaA, Hda and SeqA(16). However, it seems that cell cycle regulatory mechanism may differ under conditions supporting fast and slow growth rates. As fast growth, we assume here doubling times of 20-30 min with multiple overlapping replication rounds, whereas as slow growth rate – doubling times exceeding 70 min with single, complete chromosome duplication round. In addition to adhering to those general principles, preservation of genomic integrity during the cell cycle requires solving many particular problems by bacterial cells, for instance handling replication-transcription conflicts or coordinating DNA modification, structure and repair with chromosome duplication and segregation. Changes in growth rate and doubling time, as well as their underlying molecular mechanisms, are part of bacterial adaptation and stress responses, which are crucial for their thriving in the environment across evolutionary time scales. Uncovering changes in the replication protein interaction network may foster identification of particular mechanisms employed by bacterial cells to coordinate the cell cycle under various conditions. In this work, we took up a proteomics-based approach to test changes of replication proteins sub-network under three disparate growth conditions. We show that interactions formed by each of the eight chosen proteins change dramatically with growth conditions, except those responsible for assembly of stable complexes, like DNA polymerase III (DNA Pol III) (17), Hda-DnaN(18), NrdA-NrdB(19)and DnaB-DnaC (Fig. 1C) (20). We suggest that this remodeling may be crucial for regulation of DNA replication under different environmental conditions. Results of our screen imply also a link between DNA replication and other cellular processes, like RNA modification and degradation or synthesis of lipopolysaccharide (LPS). Moreover, they also provide alternative explanation for a few previously observed interactions made by replication proteins-encoding genes. In addition, we suggest functional relations for several unannotated genes.

**Figure 1.**
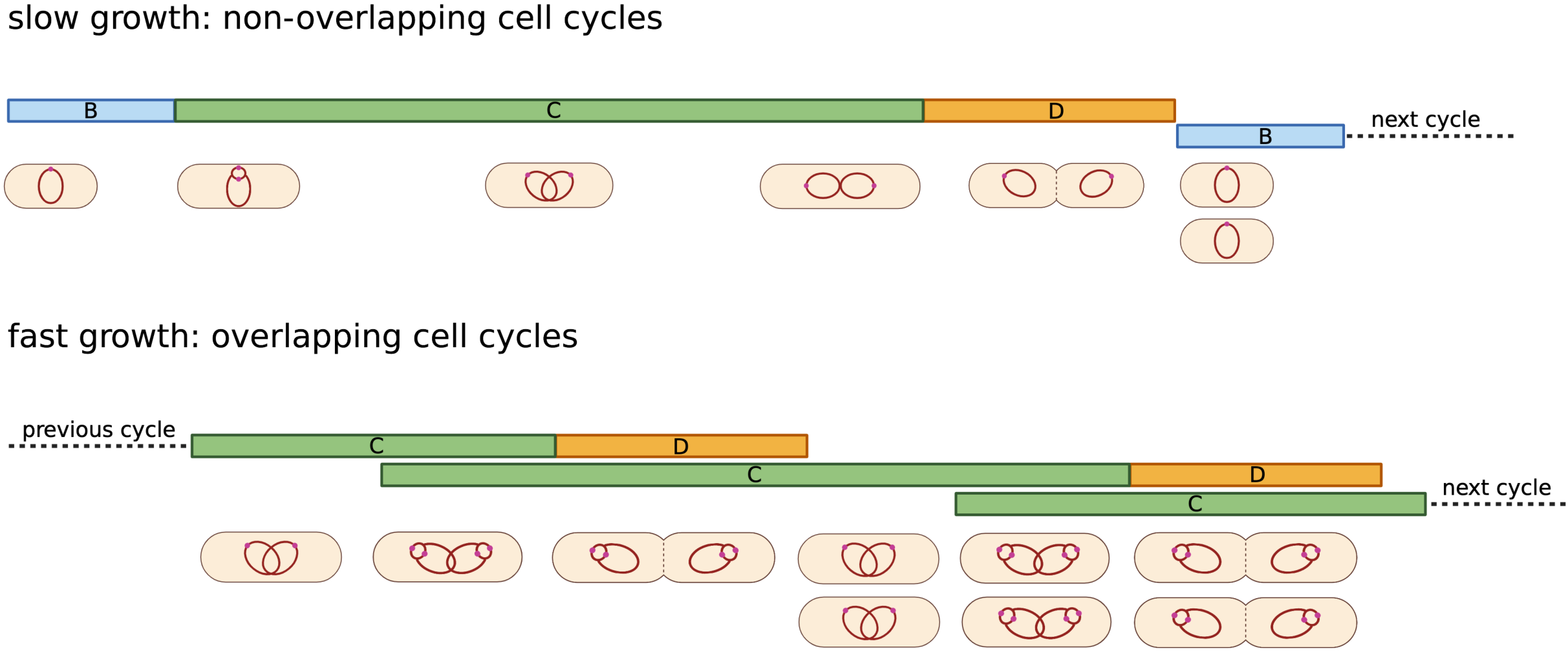
Bacterial cell cycle. A) Cell cycle stages during overlapping and non-overlapping cell cycles. B – period from cell birth to initiation of replication, C – period of time required for synthesis of daughter chromosomes, D – period required for cell division; B) DNA replication starts at a constant cell mass (volume) per replication origin irrespective of growth rate (initiation mass, invariant unit cell, IUC). Cell size increases with growth rate and overlapping replication rounds occur in fast-growing cells. Red dots represent replication origin. C) Experimentally confirmed protein-protein interactions during replication complex formation (left panel) and those recapitulated in this work (right panel)

## Materials and methods

### Strains, primers, and plasmids

List of all *E. coli* strains used in this study is shown in Supplementary table 1. Plasmids and primers used in this study are listed in Supplementary tables 2 and 3, respectively. Cloning of pUC19-pIVSK was performed using restriction-free cloning procedure, as described in (21).

### Bacterial cultures and media

LB Lennox medium (0.5% yeast extract, 1% tryptone, 0.5% NaCl) and M9 acetate medium (1x M9 minimal salts, 2 mM MgSO4, 0.1 mM CaCl2, 0.05% thiamine, 25 µg/ml uridine, 0.2% sodium acetate) were purchased from either Carl Roth GmbH or Sigma-Aldrich (now Merck). Overnight cultures were grown in LB Lennox medium. Ampicilin (Sigma-Aldrich) or kanamycin (Sigma-Aldrich) were added to the final concentration of 50 µg/ml where aplicable.

Bacterial cultures for protein complexes purification were inoculated with 20 ml of an overnight culture into 2 l of media and grown at 37°C to late exponential phase (OD_600_=0.6-1.0) (in case of LB log and M9 acetate) or to stationary phase.

### Construction of SPA-tagged *E. coli* strains and deletion mutants

All strains used for isolation of bacterial protein complexes were based on MG1655 genetic background (Supplementary table 1). SPA-tagged strains were constructed by one-step integration of linear DNA fragment containing SPA-tag sequence and kanamycin resistance cassette using λ-Red recombination method (22). DNA fragments for integration were PCR-amplified using genomic DNA of commercial SPA-tagged strain as template and primers that consist of 20nt sequences specific to SPA-tag-kan^R^ and 50nt 5’-overhangs homologous to the chromosomal regions on either side of the integration site. Deletion strains were also constructed using λ-Red recombination method (22) relying on gene replacement with kanamycin or chloramphenicol resistance cassettes. Antibiotic resistance cassettes were PCR-amplified using pKD3 or pKD13 plasmids and primers containing 50nt 5’-overhangs homologous to the region upstream and downstream of the targeted gene. PCR products were column-purified, eluted with ultra-pure distilled water and used for electroporation of MG1655 strain harboring pKD46 plasmid, expressing the λ-Red recombination proteins.

Electrocompetent cells were prepared from MG1655/pKD46 strain grown in LB+amp at 30°C to an OD_600_ ∼0.6, followed by induction with 0.15% L-arabinose and further growth for an additional one hour at 37°C. The cells were further pelleted and washed twice with ice-cold distilled water and once with ice-cold 10% glycerol. Cells were resuspended in 10 % glycerol at 100-fold concentration. For transformation, 80 µl of competent cells was mixed with ∼1 μg of PCR product. Electroporation was done with Eppendorf Eporator using 2500 V and 0,1 cm chambers. Electroporated cells were added to 1 ml LB, incubated for 3h at 37°C and plated on LB with an appropriate antibiotic. Positive transformants were PCR-verified, sequenced and subjected to FRT-FLP recombination to remove antibiotic resistance cassette, as described in (22).

### Protein complexes purification

Isolation of SPA-tagged bacterial protein complexes was performed according to protocol published by Babu and coworkers (23), with several modifications. Briefly, cell pellets harvested by centrifugation were resuspended in 20-40 ml of sonication buffer (20 mM Tris pH 7.9, 100 mM NaCl, 0.2 mM EDTA, 10% glycerol, 0.1 mM DTT, supplemented with 1 tablet of Pierce™ Protease Inhibitors (Thermo Scientific, A32965) per 50 ml of buffer) and lysed by sonication. Cell debris was removed by centrifugation at 18000 rpm for 45 min. Cleared protein lysate was incubated with 50-75 U of Viscolase nuclease (A&A Biotechnology) for 30 min on ice. After nucleic acids degradation, Triton X-100 was added to final concentration of 0.1%. The lysate was further incubated with 250 µl of prewashed in AFC buffer (10 mM Tris pH 7.9, 100 mM NaCl, 0.1% Triton X-100) Sepharose® 4B-200 (Sigma-Aldrich), for 1h at 4°C with gentle rotation. This step was performed to decrease unspecific resin binding. The lysate was cleared from Sepharose® 4B-200 and further incubated with prewashed anti-FLAG Sepharose (Biotool, B23102), for 3h at 4°C with gentle rotation. The anti-FLAG Sepharose was collected by centrifugation at 4000 rpm for 15 min and transferred into mini-spin column. The resin was washed three times with 250 µl AFC buffer and twice with 250 µl TEV cleavage buffer (50 mM Tris pH 7.9, 100 mM NaCl, 0.1% Triton X-100). Eight µl of in-house purified TEV protease (conc. ∼5 mg/ml) was added in 250 µl of TEV cleavage buffer on a column, sealed and incubated overnight at 4°C. Further, the supernatant containing cleaved proteins was collected, and CaCl_2_ was added to a final concentration of 1.5 mM. The proteins were loaded on a column with prewashed in CBB buffer (10 mM Tris pH 7.9, 100 mM NaCl, 2 mM CaCl2, 0.1% Triton X-100) Calmodulin Sepharose (GE Healthcare, 17-0529-01) and incubated for 3h at 4°C with gentle rotation.. The protein-bound resin was washed twice with 250 µl of CBB buffer and three times with 250 µl of CWB buffer (10 mM Tris pH 7.9, 100 mM NaCl, 0.1 mM CaCl2). Dried resin was stored at - 20°C prior trypsin digestion for Liquid Chromatography coupled to tandem Mass Spectrometry (LC-MS/MS).

### Identification of proteins by LC-MS/MS

Dried resin was resuspended in 50 μl of 100 mM NH_4_HCO_3_, reduced with TCEP and alkylated with iodoacetamide for 45 min at RT in dark, followed by overnight digestion with 10 ng/μl trypsin. Digestion was stopped with 5%TFA to a final concentration of 0.1%, followed by addition of acetonitrile to a final concentration of 2%. The resulting peptide mixtures were separated and measured at an online LC-MSMS setup. LC (Waters Accuity) RP-18 pre-columns (Waters), nano-HPLC RP-18 column (internal diameter: 75 μM, Waters) using an acetonitrile gradient (2%–35% ACN in 180 min) in the presence of 0.1% trifluoroacetic acid at a flow rate of 250 nl/min. The column outlet was directly coupled to the ion source of an Orbitrap Elite mass spectrometer (Thermo Scientific). Three-blank-washing runs were done between each sample to ensure the absence of cross-contamination from preceding samples. The mass spectrometer was operated in a data-dependent mode.

Analysis was performed at the Laboratory of Mass Spectrometry (IBB PAS, Warsaw). Data were analyzed using MaxQuant 1.6.3.4, referenced to *E*.*coli* proteome from UniProt database downloaded on 25.05.2020, 4391 entries. In total, 1600 proteins were identified (FDR 1%). The error ranges for the first and main searches were 20 ppm and 6 ppm, respectively, with 2 missed cleavages. Carbamidomethylation of cysteines was set as a fixed modification, and oxidation and protein N-terminal acetylation were selected as variable modifications for database searching. The minimum peptide length was set at 7 aa. Both peptide and protein identifications were filtered at a 1% false discovery rate and were thus not dependent on the peptide score. Enzyme specificity was set to trypsin, allowing cleavage of N-terminal proline. A ‘common contaminants’ database (incorporated in MaxQuant software) containing commonly occurring contaminations (keratins, trypsin etc.) was employed during MS runs. Data were deposited in Pride Repository under an entry PXD030113.

### Protein complexes - data analysis

To filter out non-specific interactants, i.e., prey proteins that are more abundant in the negative controls than in the specific experiments, statistical analyses were carried out using a custom Python script (version 3.7.6). First, based on protein intensity values, median normalization was performed for each sample to reduce inter-sample variability, followed by log10 transformation. Proteins not identified in a sample with 0 intensity value, were replaced with intensity value=1 to enable logarythmic transformation of the data.

For each prey protein record, the ratio magnitude between an experimental sample and negative control was computed and its significance was assessed using a one-way z-scorewith mean standard error computed from all proteins and samples, to increase statistical power. A protein was considered to be significantly more abundant in a specific experiment than in the negative control if the ratio magnitude between the two sets was >= 1.5, and its FDR (False Discovery Rate) adjusted-pvalue was <= 0.05 (Benjamini-Hochberg correction for multiple-testing).

Venn diagrams were made with a free online tool: http://bioinformatics.psb.ugent.be/webtools/Venn/. Protein interaction networks were analyzed and done using Cytoscape ver. 3.8.2. Functional enrichment analysis was done using a free web tool - GSEA (24).

### Chromosomal DNAreplication analysis with flow cytometry

Chromosome number measurements were performed as described in Hawkins and coworkers, with several modifications (25). Briefly, cells weregrown with aeration at 37°C until OD600=0.15 in LB medium supplemented with 0.2% glucose (fast growth) or AB medium (15.1mM (NH_4_)_2_SO_4_, 42.3 mM Na_2_HPO_4_, 22 mM KH_2_PO_4_, 51.3 mM NaCl, 0.1 mM CaCl_2_, 1 mM MgCl_2_, 0.003 mM FeCl_3_, 10 μg/ml thiamine, and 25 μg/ml uridine) supplemented with 0.4% sodium acetate (slow growth). Threeml samples were collected, treated with 150 μg/ml rifampicin, and 10 μg/ml cephalexin and incubated for 4 h at 37°C with mixing. Incubation with antibiotics results in cells containing an integral number of chromosomes, corresponding to the number of replication origins at the time of drug treatment. Subsequently, cells were harvested, washed with TBS (20 mM Tris-HCl pH 7.5, 130 mM NaCl) and fixed with cold 70% ethanol overnight or longer. Additional 3 mL sample at OD_600_=0.15 was collected without antibiotic treatment and fixed as above. After sample collection, bacterial culture was grown up to early stationary phase with OD_600_ measurements to determine doubling time of bacterial population.

Prior to flow cytometry analysis, cells were resuspended in 50 mM sodium citrate followed by RNA digestion with RNase A for 4 h. Chromosomal DNA was stained with 2 mM Sytox Green (Invitrogen) and DNA content per cell was measured with BD FACSCalibur at 488 nm Argon Ion laser. MG1655 (WT) strain grown in AB medium containing one of the following carbon sources: 0.4% sodium acetate, 0.2% glucose, 0.2% glucose + 0.5% casamino acids or in LB medium with 0.2% glucose, treated with antibiotics, fixed and stained as above was usedas a standard for each flow cytometry measurement, indicating cells containing 1/2, 2/4, 4/8 or 8/16 chromosomes, respectively.

Flow cytometry data were analyzed using FlowJo ver. 10.8.0. To determine cell volume, fixed exponentially growing cells (collected without antibiotic treatment) were micro photographed using Leica DM500 microscope. Cell length and width were measured in ImageJ and cell volume was calculated with cylinder volume equation (πr^2^h) where r – half of cell width and h – cell length.

### Figures

All figures from the manuscript were prepared using https://biorender.com

## Results

### Experimental setup and data analysis of the replication proteins interaction network and its dynamics under different bacterial growth conditions

The rationale behind our experimental plan was that protein modules responsible for chromosome duplication may undergo substantial remodeling as conditions change, underlying growth rate and/or condition-dependent control of DNA replication. Therefore, we aimed to investigate the composition of protein complexes formed by the main DNA replication regulators (DnaA, DiaA, Hda, SeqA)(16) and key replication complex components (DnaB, DnaG, HolD)(17) in chosen, disparate conditions. In our analysis, we included also NrdB, a ribonuclotide reductase (RNR) subunit, the key enzyme producing deoxyribonucleotides. Experimental evidence suggests that RNR may be associated with the replication complex (26), with its activity dictating the length of the C period (27). We used AP-MS according to the protocol published previously by Butland and coworkers (23, 28) to assess protein-protein interactions of selected replication proteins. Replication machinery interactome was probed during fast bacterial growth in rich, undefined medium (LB) and in defined synthetic medium, supporting slow growth rate, with acetate as a sole carbon source. In both cases, protein complexes were investigated in the late exponential phase (OD_600_ ∼ 0.6-1.0) and for LB-grown cultures also upon cell culture entry to stationary phase. This way, we could subsequently compare changes within the replication proteins PPI network between fast and slow growth conditions, and after bacterial growth had ceased. For each protein queried, we prepared a strain producing its SPA-tagged version. We constructed C-terminally tagged translational fusions of the cognate genes, under control of their native promoters, in the MG1655 genetic background. Using such a setup we wanted to ensure near-endogenous levels of the replication proteins used as baits in our experiment, to minimize spurious interactions. Following isolation by tandem AP, composition of proteins co-purified with the baits was assessed by mass spectrometry and analyzed quantitatively using MaxQuant (29). A critical issue in AP-MS experiments is elimination and identification of protein impurities that interact either with resins or bound proteins unspecifically and obscure subsequent identification of true interactants. On the other hand, not every protein forming an interaction with the resins or SPA-tag itself should be automatically accounted for as a false-positive when analyzing samples containing tagged baits and therefore protein enrichment should be calculated in such cases. To tackle this problem, we performed two types of control experiments for each growth condition tested, which further allowed to set threshold criteria for unspecific protein binding and filter interactions characterized by substantial prey enrichment and low False Discovery Rate (FDR adjusted p-values < 0.01). Controls were composed of an untagged MG1655 strain and an MG1655 strain expressing a SPA-tagged fluorescent Venus protein, both grown under identical conditions to those used for the strains expressing the tagged bait proteins. In the first case, the control experiment enabled correction for proteins attaching unspecifically to the resins, in the second – for proteins binding to a random SPA-tagged protein or SPA-tag itself. The use of the two control types delimits abundance range of a protein that binds unspecifically, dependent on resin occupancy by a bait protein. Specifically, we made a presumption that the level of proteins binding unspecifically to the resin will be lower when the amount of bait protein and its interactants is high and thus, the resin beads are more occupied. The differences between resin occupancy among different bait proteins may result from different native protein expression levels as well as various tag surface exposition on the natively folded proteins. Considering reproducibility of the results, for further statistical analysis we took into account only protein hits that were identified in each of three sample replicates (Supplementary figure 1A-H and Supplementary file 1). However, we performed statistic tests for every protein hit that appeared in at least one replicate of the control samples. Statistic tests were performed separately for each control. We further considered as hits only the interactions that were statistically significant with respect to both control types as well as preys that did not appear in any of the controls. In similar proteomics analysis published previously, authors considered as potential hits also the proteins that appeared in both three and at least two sample replicates. We did not perform statistical analysis for the latter cases, but we selected those proteins which appear in at least two sample replicates but are not present in either of the controls and listed them in the Supplementary file 2. Moreover, we did not perform any manual curation of the data, leaving also all recovered ribosomal proteins in our dataset. Although ribosomal proteins constitute frequent contaminants in AP-MS experiments, their profile changed significantly for each protein tested and for HolD, DiaA, NrdB and Hda the fraction of ribosomal protein preys was much less abundant than for DnaA, SeqA, DnaB and DnaG. Therefore, some of the co-purified ribosomal components may represent true interactants. For instance, such direct interaction has been previously reported between DnaA and the ribosomal protein L2 (30).

### Screen results recapitulate well-known replication proteins contacts and provide possible explanations for several previously observed genetic interactions

As expected, results of the screen confirmed well-known connections between complex components of several replication proteins, like these formed by DNA polymerase III subunits, ribonucleotide reductase complex or between Hda and β sliding clamp of DNA polymerase III (Fig. 1C). These are stable complexes that were isolated under all tested conditions. Our results also recapitulated the interactions described previously as spatiotemporally-regulated during the cell cycle (DnaB-DnaC, DiaA-DnaA, HolD-Ssb, topoisomerase III-HolD) (Fig. 1C, 2) (17, 31, 32) or performing special function, i.e. replication through highly transcribed regions (DnaB-Rep)(33) (Fig. 2). These results confirm that the approach we used accurately identifies different types of complexes formed by the selected replication proteins. Moreover, our data also corroborates some of the experimentally supported interactions present in the STRING database (Fig. 2). Though, the majority of the identified interactions have not been reported in the STRING database before (Supplementary file 3). On the other hand, our screen has not captured all of the experimentally confirmed interactions found in the STRING database. This is however not surprising as most of the missing interactions were previously identified based on assays consisting of purified proteins or the yeast two-hybrid system (Supplementary table 4). In those assays, both protein partners are provided at relatively high amounts at the same time and space. Those conditions may not be recapitulated under bacterial growth conditions used in our study and instead those protein assemblies may be specific to certain physiological sates of bacterial cells or represent less prominent cellular protein complexes.

**Figure 2.**
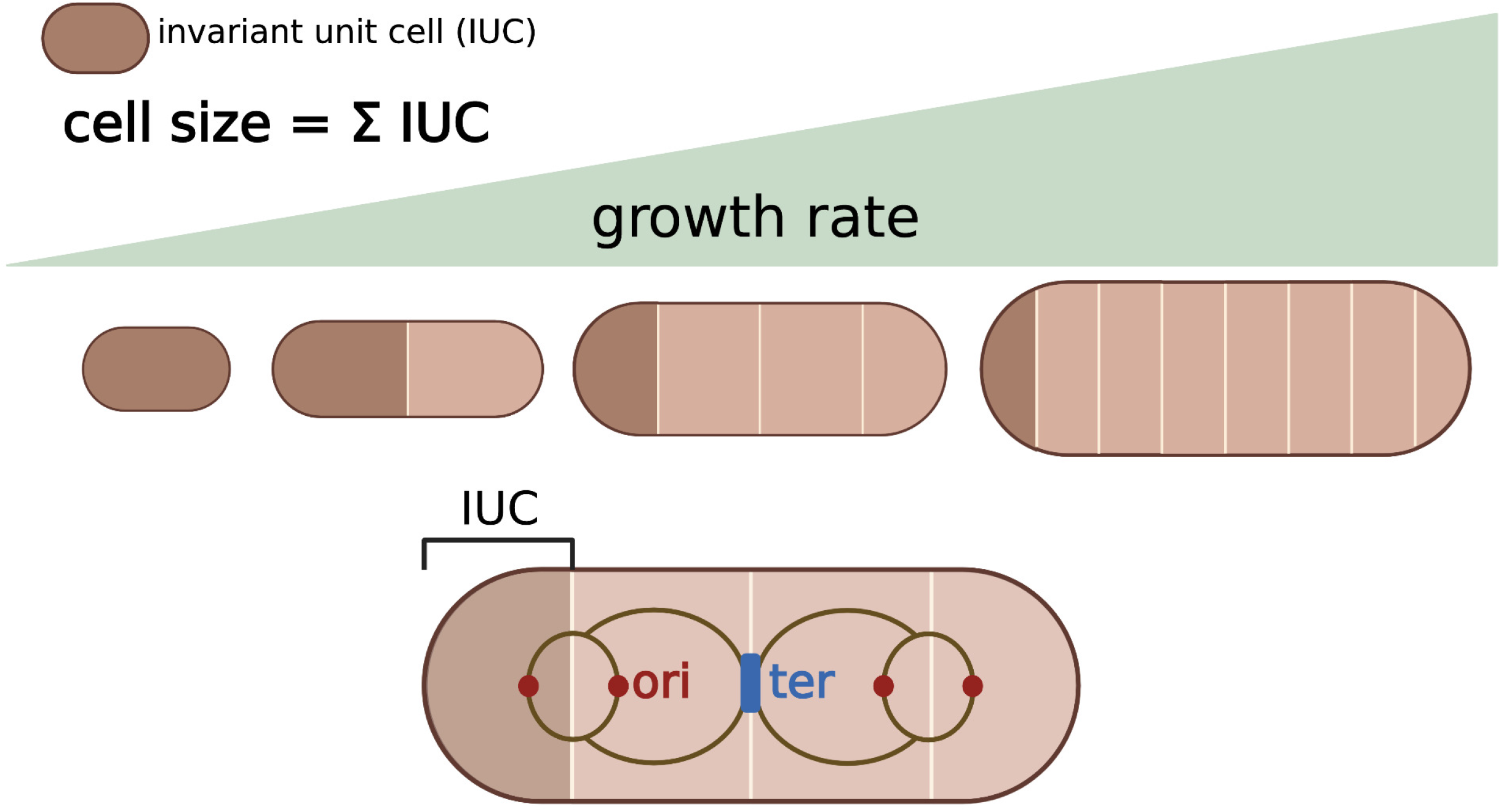
Experimentally confirmed PPIs present in the STRING database for the eight replication proteins analyzed in this work. Baits used in the screen are marked with octagons. PPIs confirmed in this worked are presented with green edges. Proteins identified in the screen as interactants of other replication proteins than their partner in the STRING database are highlighted in bold.

Interestingly, among significantly enriched preys, we have found examples of proteins which had been previously implicated in an interplay with the *E. coli* replication machinery by genetic screens (Supplementary file 2, Fig. 3). Namely, SspA protein was isolated as an interactant of HolD, whereas RlmE was found in complex with NrdB. Previously, a deletion of *sspA* has been shown to suppress growth defect of a *ΔholD* strain (34). SspA is a transcription factor required for stress and starvation responses, therefore the mechanism of suppression was suggested to be indirect and operate through changes in gene expression in a *ΔsspA* strain (35). Another study revealed that SspA may in fact be needed to solve transcription-replication conflicts (36). RlmE, on the other hand, is a 23S rRNA methyltransferase responsible for methylation of the 2’-O ribose position at the conserved U2552 nucleotide. Growth of *ΔrlmE* strains was found sensitive to hydroxyurea, a well-known inhibitor of ribonucleotide reductase (37). This sensitivity was attributed to increased membrane stress of cells devoid of RlmE, due to perturbed translation and the resultant elevated incorporation of misfolded proteins to their cell membrane. However, in light of our results, in both cases, the observed genetic interactions may have more complex cause and arise from direct protein-protein contacts of RlmE and SspA with the replication machinery. Those results show that the data produced by our screen provide meaningful results, supporting reliability of previously uncharacterized protein-protein interactions identified in this work.

**Figure 3.**
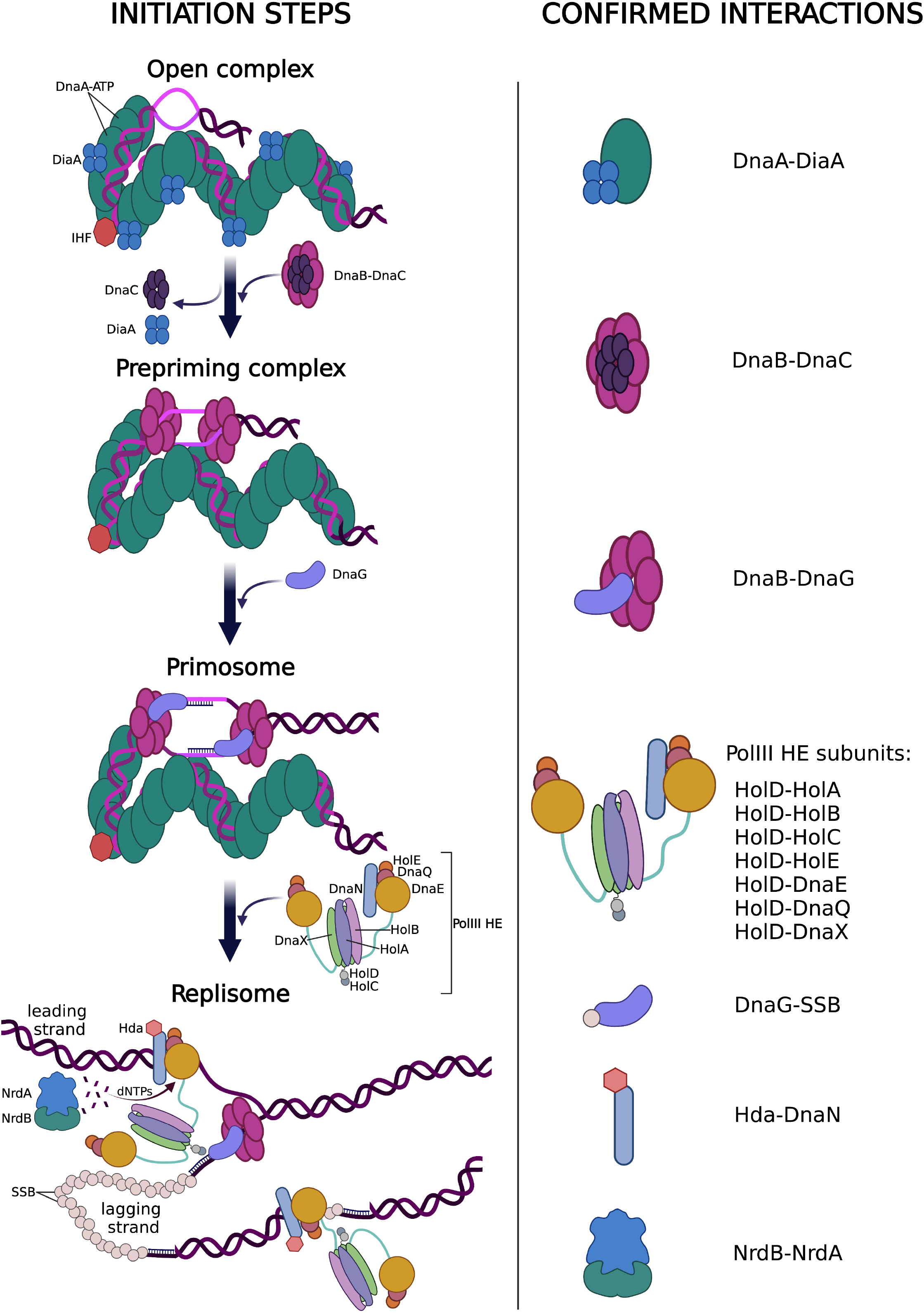
Condition-dependent protein-protein interactions formed by the eight *E. coli* replication proteins selected for this study. Briefly, protein complexes were analyzed by AP-MS. Chromosomally expressed DnaA, DiaA, Hda, SeqA, DnaB, DnaG, HolD, NrdB were C-terminally tagged with SPA-tag in MG1655 genetic background. Bacteria were grown either to middle/late exponential phase in LB medium or M9+acetate as a carbon source, or to stationary phase in LB. Bacterial cells were lysed and protein complexes were subjected to sequential affinity purification according to the protocol published in (23), with minor modifications. Isolated protein complexes were digested with trypsin and their components were identified using LC-MS/MS. Proteins were quantified using MaxQuant(29). Threshold for unspecific interactions was delimited according to results of control experiments with untagged MG1655 strain or the strain expressing moderate levels of SPA-tagged mVenus protein. Graphs present preys isolated with each of the bait proteins under all tested conditions. Edges are color-coded according growth conditions where interaction was identified and edge width is proportional to prey enrichment. Only interactions with corrected p-value < 0.01 were shown. Preys were grouped into functional classes, listed in the figure legend.

### Interaction networks are specific for each replication protein and bacterial growth condition

Results of the PPIs analysis revealed that each of the queried replication proteins forms a unique constellation of contacts with the rest of the proteome (Fig. 3, Supplementary file 4). For simplicity, preys identified in the screen were classified into arbitrary single functional categories, although some proteins could be ascribed to more than one cellular function. Nevertheless, as described in the preceding paragraphs, our screen confirmed previously found interactions among the replication proteins. In AP-MS experiments, it is possible that the identified preys do not form a direct contact with the bait, but rather with some of true primary interactants. Those preys may still be a part of a larger complex formed by the bait or comprise alternative complexes formed by the interactant. However, a small overlap between sets of preys co-purified with baits that interact with each other suggests that the selected preys, at least in part, represent direct interactions (Fig. 4). Interestingly, DnaA regulators Hda and DiaA formed a small number of interactions, whereas proteins associated with replication forks progression (HolD, NrdB, DnaB, SeqA) and DnaA formed large but distinct networks. For instance, HolD displayed many statistically significant connections with various metabolic proteins but none with the ribosomal ones, contrary to SeqA, which interacted with only two metabolic enzymes. One of them was polyphosphate kinase (Ppk), being also a component of *E. coli* RNA degradosome (38), as described in more detail in the next sections, could also be classified as a RNA modification machinery member. In general, similarity of the PPI network (percentage of the same preys) was the highest for DnaA and SeqA and high also for DnaB and SeqA, although SeqA has not been found to interact directly with either DnaA or DnaB (Fig. 4). In both cases, however, high similarity stems from a large number of significantly enriched ribosomal proteins common between the three baits which are likely in part false-positives. Though, unlike DnaA, DnaB shares with SeqA ribosomal protein interactants L16 (RplP) and L23 (RplW) (LB, log phase) (Fig. 3) that were absent from all controls and thus are considered as hits. It is also worth noting that the two ribosomal proteins were identified as a part of the DnaG protein complex. Primase by itself co-purified with SeqA (LB, stationary phase; Fig. 3), indicating that the two proteins may interact. Such interaction hasn’t been described before for the *E. coli* replication complex and its role remains unknown.

**Figure 4.**
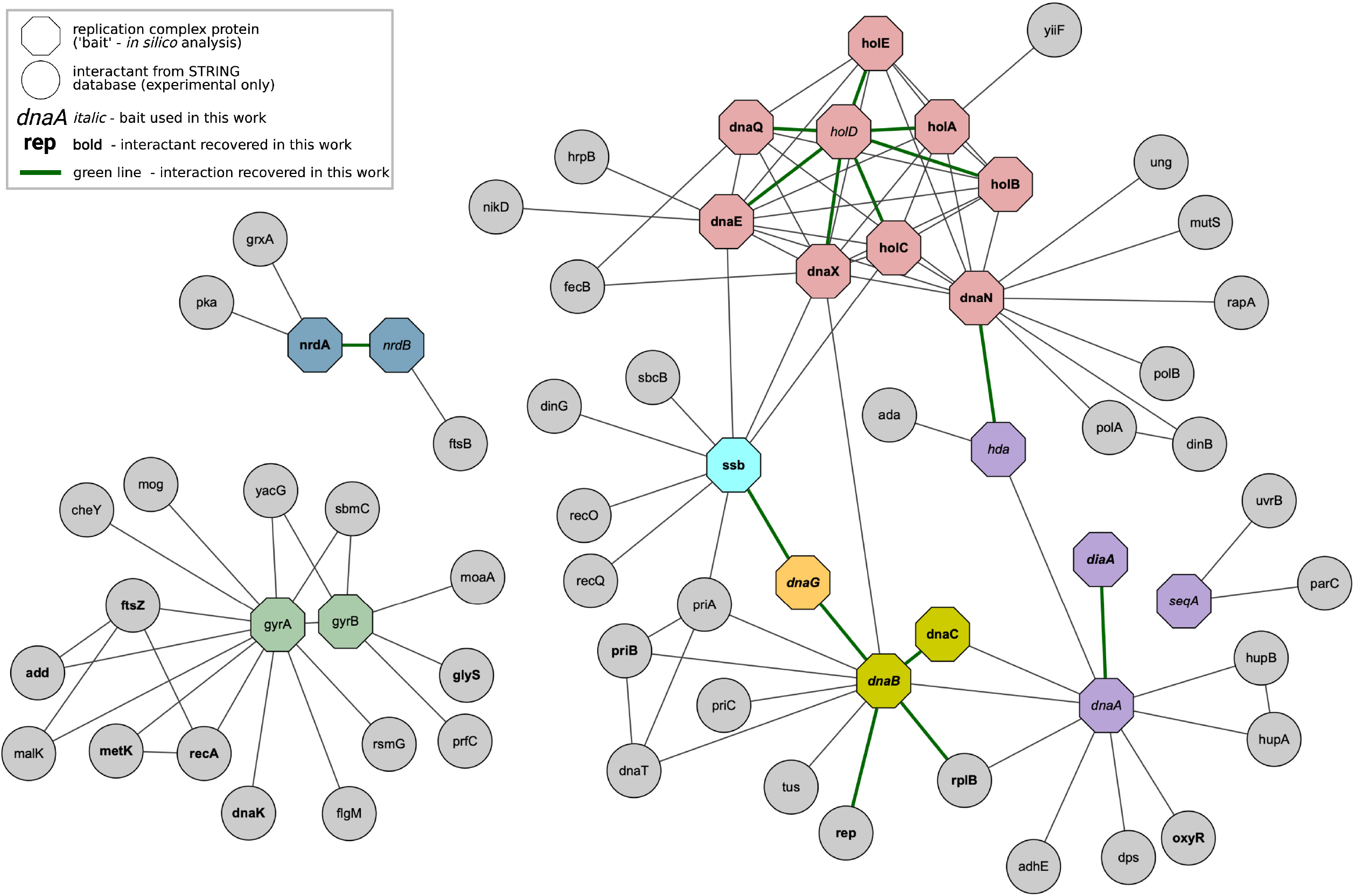
Pairwise comparison of uncovered interaction profiles of 8 baits used in this work. Similarity matrices were made based on comparison of interaction profiles between each of 8 bait proteins used in PPI screen (‘each with each’ comparison). Percent of common interactants between compared bait pairs was calculated and subsequently weighted average was calculated of the number of all interactants identified in our PPI screen.

Importantly, the set of interactants for every bait used in our screen changed drastically with growth conditions (Fig. 5A). In general, the highest number of statistically significant interactions was observed in samples obtained from bacteria during their exponential growth in rich medium, whereas the smallest – in samples grown in minimal medium with acetate (Fig. 5A). This difference in statistically significant prey number can only partially be attributed to bait abundance in samples from different growth conditions (Fig. 5B). At the same time, stable complexes, like DNA Pol III, Hda-DnaN or DnaB-DnaC were isolated under all conditions, showing that the procedure provides meaningful results irrespective of the culture growth conditions. Those results suggest also that PPI modules comprising the tested replication proteins undergo significant reorganization as environmental conditions change.

**Figure 5.**
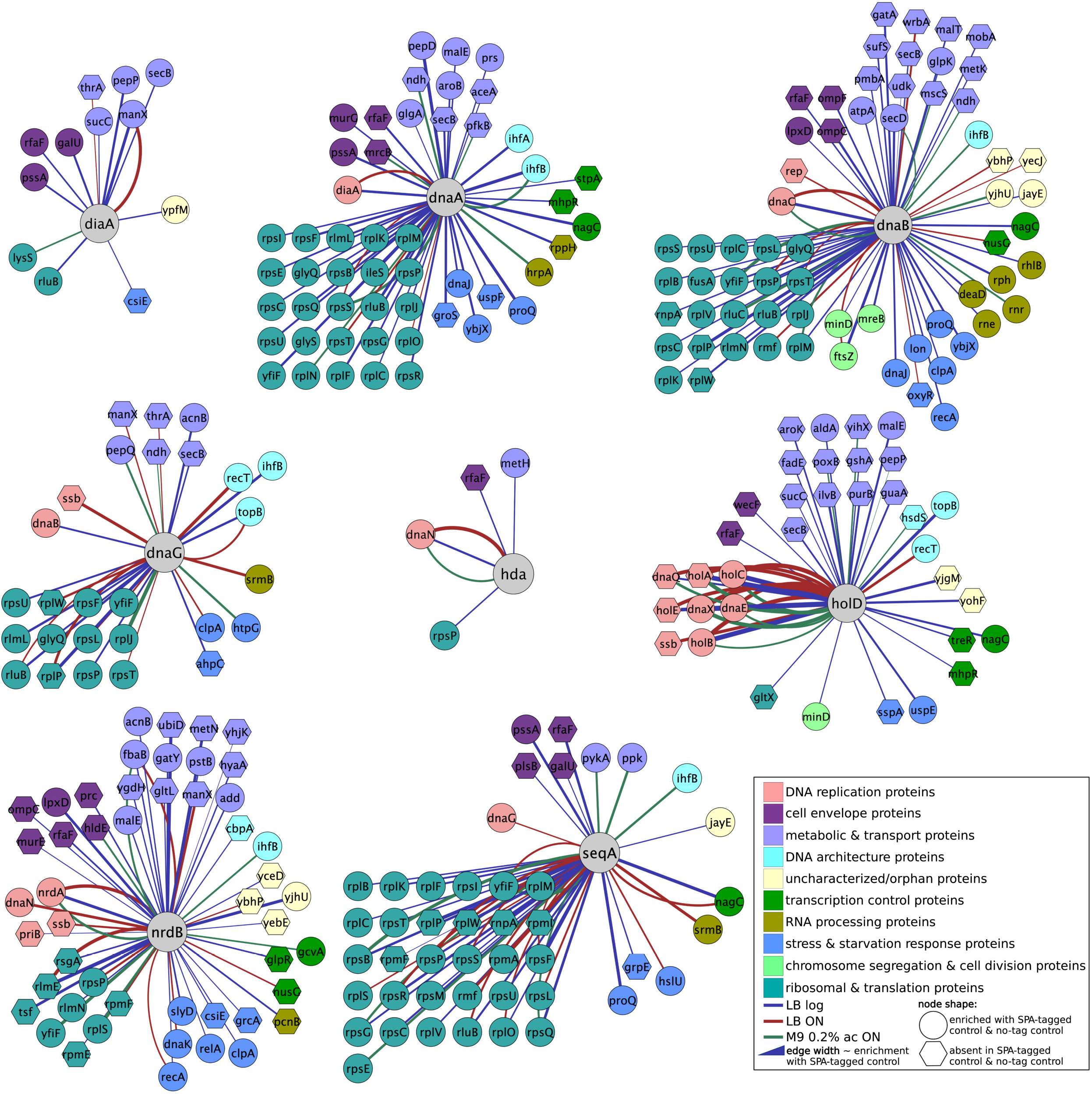
Venn diagrams showing similarity of interaction profiles for each bait protein used in this study under different conditions (upper panel). Diagrams presenting bait abundancies in the AP-MS experiments, under different growth conditions (showed as log10 value of bait intensity) (lower panel). Protein intensities were calculated using MaxQuant.

### Functional enrichment analysis of PPIs formed by different replication proteins

Functional enrichment analysis of PPIs formed by six of the tested replication proteins in all tested conditions revealed several interesting features (Fig. 6; Supplementary file 6). The number of interactants isolated with DiaA and Hda was too low to perform this kind of analysis. As mentioned earlier, HolD protein sub-network was enriched in metabolic enzymes (Fig. 6). Two of them (PurB and GuaA) are members of purine nucleotide synthesis pathways, suggesting possible coordination between nucleotides pool and DNA replication. Interestingly, four proteins were enriched in interactants with RNA-binding function (DnaB, DnaG, DnaA, SeqA). In addition, DnaB was enriched in preys with ribonuclease activity (Fig. 6). Closer examination of the data revealed that two of the above mentioned proteins (DnaB and SeqA) were co-purified with components of the *E. coli* degradosome (Fig. 3)(38). Specifically, DnaB pulled-down RNase E (Rne), RhlB and also Ppk (p-value = 0.01) in the samples from the exponential growth in LB, whereas SeqA was isolated with ppk. SeqA also pulled-down SrmB, which interacts directly with RNase E, in samples isolated from LB overnight culture. SrmB was also found as a prey in the case of DnaG. As mentioned earlier, DnaG and SeqA were found to co-purify with each other. However, statistical significance was found in the case of SeqA bait and DnaG prey in the samples isolated from LB overnight culture, whereas in the reversed experiment SeqA co-immunoprecipitated significantly with DnaG in the samples from exponential phase (p-value = 0.01). Those results suggest that DnaB and SeqA may both interact with the RNA degradosome, while SeqA interacts also with DnaG. All those interactions have not been described before.

**Figure 6.**
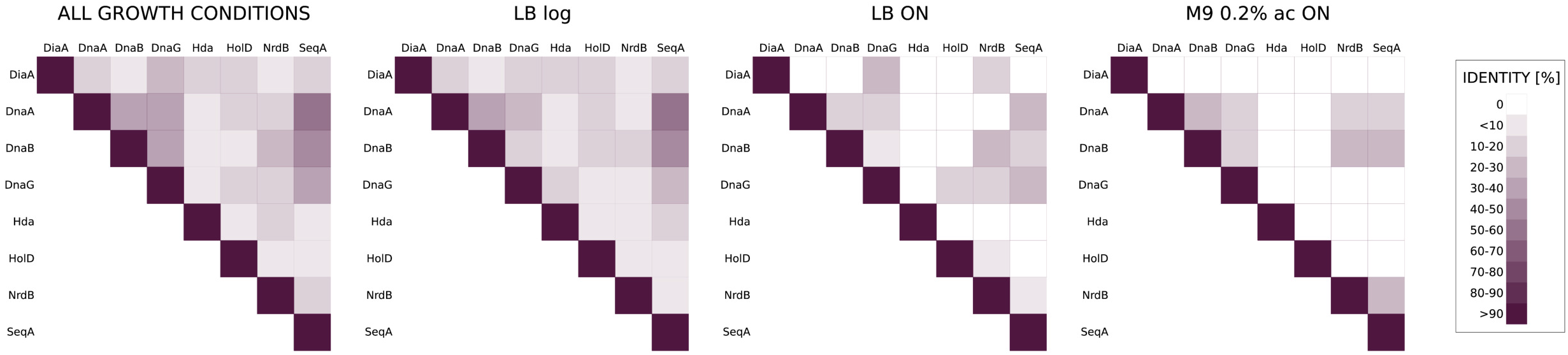
Functional enrichment analysis of interactions formed by six of the eight bait proteins used in this work. Analysis was performed with web tool GSEA, described in(24). The entire statistically significant prey set (p-value < 0.01) of each bait protein, obtained from each of the tested growth conditions, was used as input for analysis.

Interestingly, our analysis recovered many connections between protein involved in cell envelope biogenesis and the replication proteins. Namely, DnaA was enriched in peptidoglycan synthesis proteins (MurG, MrcB) in the functional enrichment analysis. All other replication proteins tested have also shown conspicuous number of interactions with enzymes related to synthesis of the cell envelope precursors or cell envelope integral proteins, although GO term enrichment analysis did not show statistical significance in those cases. In particular, proteins engaged in the lipopolysacharide (LPS) synthesis (39) were purified with NrdB (RfaF, HldE, LpxD), DnaB (LpxD and RfaF), HolD (RfaF, WecF) and SeqA (RfaF). While RfaF was present as prey in many experimental samples (highly-enriched with DnaB and SeqA baits), it is worth mentioning that it was absent from all but one of the eighteen control samples (from all conditions), where it was identified in minute amounts. SeqA was also found to co-immunoprecipitate with PlsB, an enzyme engaged in a membrane phosphlipids synthesis. Intriguingly, we have also found GalU, previously implicated in cell division control through its product (40), as prey of DiaA and to a lesser extent – SeqA. Those interactions seemed very specific as GalU was also present in very small amounts in only one of 18 control samples. A connection between cell envelope biogenesis and DNA replication has been suggested by many previous studies which we discuss in more detail below.

NrdB subunit of RNR, as expected, was enriched in DNA replication proteins – Ssb, PriB and DNA Pol III subunit DnaN. The latter may be the RNR contact point within the replication complex.

### Interaction networks components have functional impact on replication timing in the E. coli cell cycle

While cell cycle stages of fast-growing bacteria occur simultaneously, it is crucial for the cell to provide precise timing of the replication initiation in each cell cycle as well as to couple each replication initiation with corresponding cell division according to ‘one initiation of replication per cell cycle’ rule (12) (Fig. 1A). The main conclusion of numerous research on the interplay between bacterial growth and cell cycle is that cells maintain growth rate-dependent size homeostasis and that the relationship between cell size, replication timing and thus the DNA content in the cell is precisely regulated (14, 15) (Fig. 1B). As a consequence, each change in DNA content under specific growth rate should be accompanied with changes in growth rate and/or cell size. If these relationships are disturbed, bacterial population exhibit deregulations in precise replication timing, namely the initiation might occur too early or too late in the cell cycle. Moreover, problems in proper replication timing may also lead to even more serious effect – origins present in the cell may not fire simultaneously which results in uneven distribution of genetic material to progeny cells.

Consequently, chromosome copy number measurement along with size and growth rate calculations can be used as a way to determine the replication timing in exponentially growing bacterial population. In this approach, centered on flow cytometry, chromosomal content in bacterial cells is assessed after the so-called replication runout performed with rifampicin and cephalexin treatment (25). Upon inhibition of the initiation of new rounds of chromosome replication and cell division with those antibiotics bacterial cells remain with the chromosome number corresponding to the number of origins present at the time of antibiotics addition. DNA content can be then measured with flow cytometry and correlated with microscopically-determined cell size and growth rate of bacterial population derived from OD measurements (25, 41). Since the chromosome number present in *E. coli* cells growing in defined media has been well established before (41), the wild type strain can serve as a standard to compare with mutant strains.

To determine whether replication protein interactants recovered in this work can be functionally involved in replication regulation during the cell cycle, we constructed deletion mutants of chosen genes encoding those proteins and subjected them to the analysis described in the preceding paragraph. We chose genes from two different functional groups, namely: *rlmE*, gene encoding 23S rRNA methyltransferase found in complex with NrdB and *rfaD*, gene encoding ADP-L-glycero-D-mannoheptose 6-epimerase found in complex with SeqA (p-value = 0.01). RfaD is engaged in LPS synthesis and related in several ways to another significant interactant identified in our screen – RfaF. Namely, RfaD is responsible for the synthesis of substrate metabolite that is directly used for the reaction mediated by RfaF. Besides, rfaD and rfaF genes are transcriptionally co-regulated within a single operon. We chose to analyse RfaD, because the type of analysis described above was not possible with *ΔrfaF* mutant, since it does not grow in minimal medium with acetate as carbon source (data not shown).

Deletion mutants were grown in two different conditions, providing fast and slow growth rate and corresponding to the conditions used for the PPI screen. Cells in early logarithmic growth phase (OD600=0.15) were subjected to replication runout and total chromosome copy number was measured using flow cytometry. Results for *rlmE* and *rfaD* deletion mutants are presented in Figures 7 and 8, respectively. *E. coli* MG1655 (WT) strain cultured in different media served as a reference strain to make a standard curve depicting the relationship between chromosome number and cell volume as well as between chromosome number and doubling time (supplementary fig. 2 and supplementary file 7).

**Figure 7.**
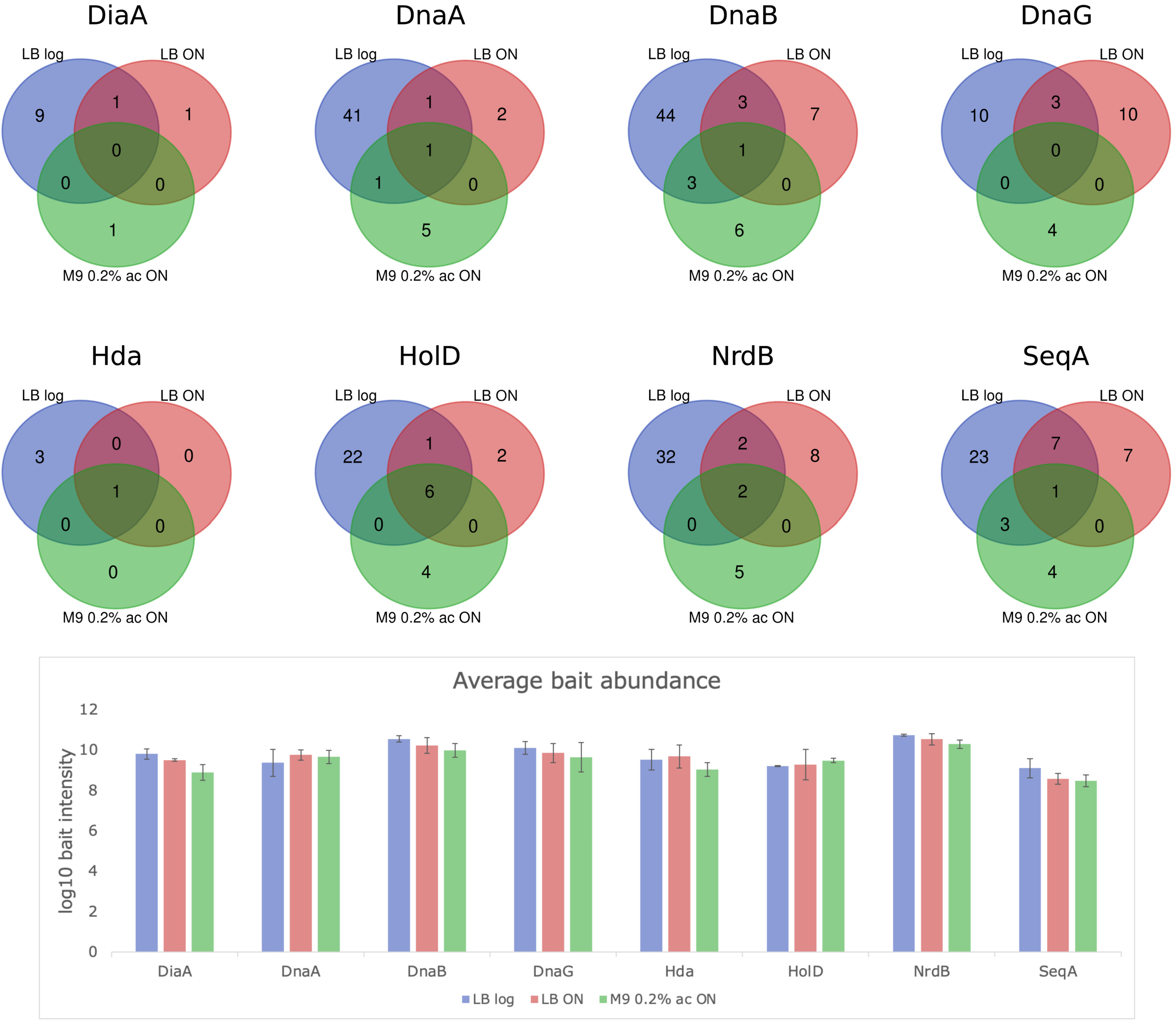
Cell cycle parameters of the *ΔrlmE* strain. (A) Upper panel - flow cytometry histograms presenting cell populations containing particular chromosome number, lower panel – microphotographs of representative cells from the population. Chromosome number, present in *E. coli* cells grown in LB+glucose or AB medium + acetate, was calculated after replication run-out, Sytox Green staining and flow cytometry analysis. (B) Graphs show an interplay between chromosome number and cell volume (left) or doublings per hour (right). Cell volume was measured using ImageJ software after collecting cell images with light microscope and phase contrast. OD_600_ measurement of exponentially growing cultures served as basis of growth rate calculation. WT *E. coli* strain cell cyle parameters were set up as a standard curve to which mutant strains can be compared. *p-value<0.05 (t test) *** p-value<0.001 (t test)

**Figure 8.**
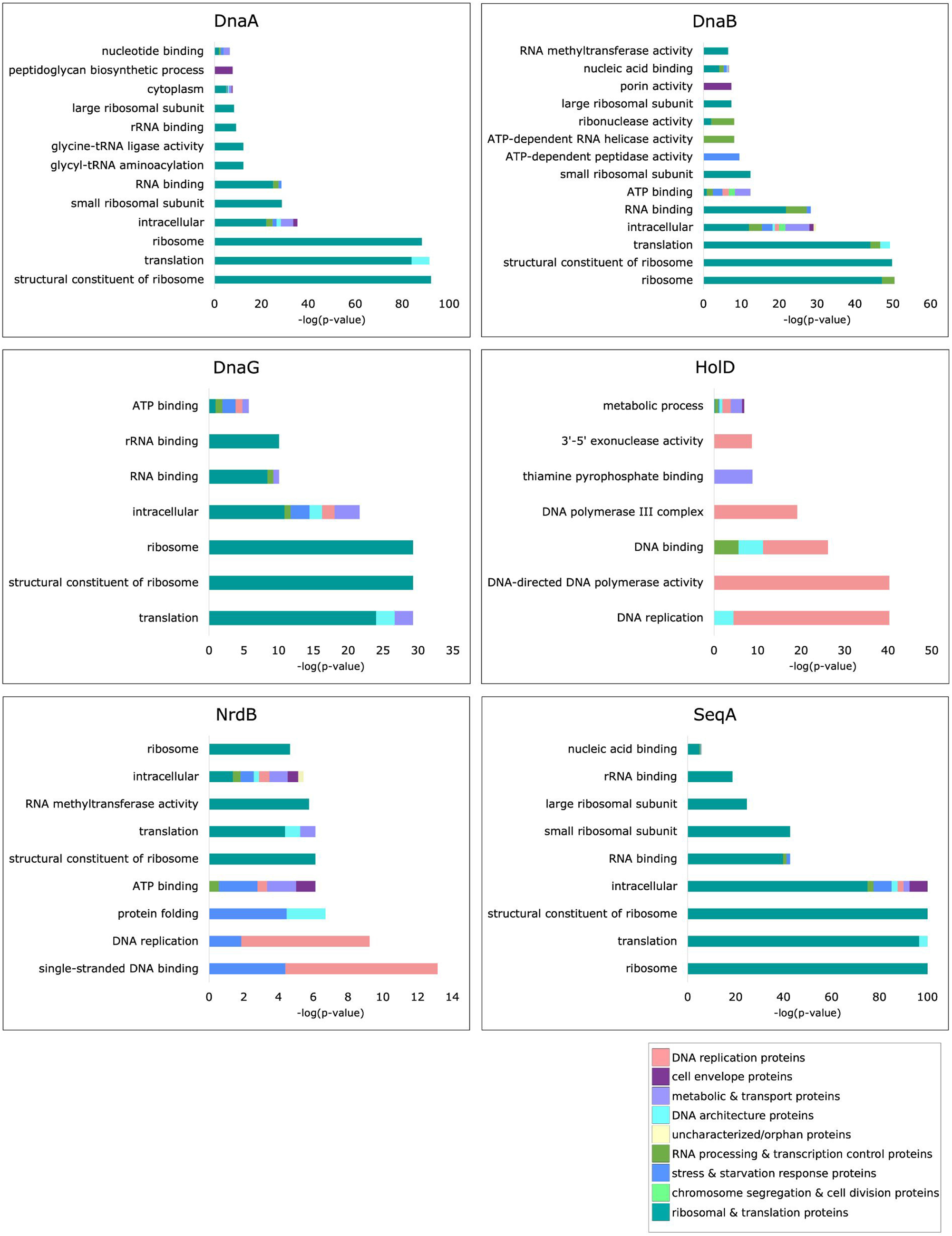
Cell cycle parameters of the *ΔrfaD* strain. (A) Upper panel - flow cytometry histograms presenting populations containing particular chromosome number, lower panel – microphotographs of representative cells from the population. Chromosome number, present in *E. coli* cells grown in LB+glucose or AB medium + acetate, was calculated after replication run-out, Sytox Green staining and flow cytometry analysis. (B) Graphs show an interplay between chromosome number and cell volume (left) or doublings per hour (right). Cell volume was measured using ImageJ software after collecting cell images with light microscope and phase contrast. OD_600_ measurement of exponentially growing cultures served as basis of growth rate calculation. WT *E. coli* strain cell cyle parameters were set up as a standard curve to which mutant strains can be compared. *p-value<0.05 (t test) *** p-value<0.001 (t test)

Our results suggest that both deletion strains exhibit disturbances in proper replication timing, especially under the conditions supporting fast bacterial growth rate. The majority of cells within the populations of Δ*rlmE* and Δ*rfaD* mutants grown in LB medium supplemented with 0.2% glucose contained 4 and 8 chromosomes (less than WT cells that had 8 and 16 chromosomes under these growth conditions), both mutant populations contained a fraction of cells where asynchronous replication events occurred. Slowly growing Δ*rlmE* cells (minimal medium +acetate) showed no differences in chromosome number comparing to WT strain, whereas *ΔrfaD* population encompassed a small fraction of cells with higher DNA content.

Measurements of cell size and doubling time revealed that Δ*rlmE* strain grew slower than WT in rich medium, with elongated cells and bigger cell volume (supplementary table 4). Δ*rfaD* deletion mutant, in turn, exhibited slightly smaller cell volume in both growth conditions whereas doubling time was much longer than in comparison to the WT strain only in the minimal medium (Supplementary table 5). These values plotted on standard curve graphs suggested that the relationship between cell volume and DNA content is significantly disturbed in Δ*rlmE* deletion mutant under the conditions supporting fast growth rate. The interplay between chromosome number and cell cycle duration (expressed as cell doublings per hour) is also deregulated in fast-growing Δ*rlmE* strain, but also in slowly growing rfaD deletion mutant. Those results suggest that RlmE and RfaD indeed have an impact on cell cycle regulation and their relation with RNR and SeqA remains to be established.

## Discussion

Protein-protein interactions are crucial for execution and regulation of all cellular functions. Cell cycle is a particular example of process regulated by spatiotemporally resolved protein complexes. It is because complexes composition has to change at different cell cycle stages to enable performance of different biochemical functions required at different steps. At the same time, coordination of genome duplication with other DNA transactions, like chromosome segregation and transcription, has to be ensured. The latter may also mean that different solutions are needed under different gene expression programs. Some of the protein complexes operating at chromosome replication constitute stable multisubunit machines, others rely on transient protein-protein interactions. For regulation of molecular processes, it is important not only whether a complex is formed or not but also – if its more or less abundant and how this affects the possibility of formation of alternative complexes. Thus, to fully understand regulation of DNA replication in bacteria at different environmental regimes, it is important to qualitatively and quantitatively assess the dynamics protein interaction network under various conditions. The major driving force of changes in protein complexes and entire modules are alterations in protein abundance and hence - abundance of the alternative interaction partners of hub proteins. However, whereas expression of genes encoding components of stable complexes is usually coordinated, it is not necessarily true for transient interactions. Thus, transient interactions may be more difficult to predict from gene expression data alone. Some of the mechanisms responsible for such interactions are post-translational protein modifications. Already existing *E. coli* protein-protein interaction networks are usually static and do not provide any information on how composition of protein complexes and protein modules changes under different conditions. The same is true for well-established components of the *E. coli* replication and pre-replication complexes. The crucial interactions (for instance DnaA-Hda, DnaA-DiaA, Hda-DnaN) (Fig. 1C) are characterized but it is not known whether they all play a role under various conditions and if additional factors may be operational. In this work we analyzed the PPI network of the key proteins involved in DNA replication in *E. coli*. For this proof of concept study we chose label-free quantitative proteomics approach. We have shown that the number of interactions formed by each tested replication protein changes significantly with growth conditions. Those results corroborate the hypothesis that the network of replication machinery interactions changes to accommodate variations in DNA replication itself and other processes occurring on DNA (Fig. 3). We demonstrate that some interactions appear and dissolve under different conditions. Absent interactions in our results do not necessarily mean that they do not exist under particular conditions at all, but that they are less abundant or less stable. Our results also suggest that the high-throughput studies on PPIs, performed on stationary phase cells provide a fragmentary picture on the interaction network formed by the replication proteins.

Our screen performed on exponential cultures revealed several interesting interactions. First of all, DnaB and also SeqA were co-purified with components of the RNA degradosome (Fig. 3, Supplementary file 2). While, to our knowledge, no evidence on cooperation between the replication machinery and RNA degradosome exists so far in *E. coli*, it has been found that the function of RNA surveillance machinery is required for stability of human mitochondrial genome (42). Activity of human RNA degradosome is required for prevention of harmful R-loop formation in the mitochondrial genome. As R-loops impede replication fork progression in *E. coli* (43), it is possible that such interaction would foster R-loop removal and ensure processivity of the replisome. Similarly, stress response transcription factor SspA has been previously suggested to play a role in resolving transcription-replication conflicts(36). Our results indicate that a direct protein-protein interaction of SspA with HolD, or indirect – with other HolD interactant – may play a role in this SspA function (Fig. 3, Supplementary file 2). One of the overrepresented functional protein categories among the preys, belonging also to RNA-modifying enzymes, were RNA methyltranferases (RlmE, RlmL) (Fig 6). We paid particular attention to one of them – RlmE, due to high specificity of the interaction, its condition-dependence and previously described phenotypic relation to the cognate replication protein bait – RNR. We analyzed cell cycle parameters in *E. coli* cells devoid of the *rlmE* gene and found that its absence affected replication synchrony and the correlation between chromosome number, cell volume and growth rate under fast growth conditions (Fig. 7).

Secondly, our screen revealed many interactions of replication proteins with the cell envelope (inner membrane-peptidoglycan-outer membrane) proteins or enzymes directly engaged in the synthesis of cell envelope precursors (Fig. 3, Supplementary file 2). Some connections of the replication regulators DnaA and SeqA with the cell membrane and outer membrane have been suggested before, as summarized in Fig. 9A. Namely, an interaction between acidic phospolipids and DnaA has been shown to enhance the exchange rate of ATP/ADP nucleotide bound by DnaA (44) and inhibit DnaA binding to *oriC* (45). In addition, a depletion of PgsA, an enzyme necessary for synthesis of acidic phospholipids, has been demonstrated to result in cell cycle arrest (46). The mechanism of this arrest is still not fully understood but a large body of evidence suggests that it is dependent on a membrane-binding region of DnaA (47, 48). Moreover, the negative replication regulator SeqA has been found to be responsible for hemimethylated *oriC* binding by membrane fractions (49) and be by itself associated with the cell inner membrane (50). What is more, *ΔseqA* mutants have been shown to exhibit increased LPS phosporylation, contributing to elevated ori/ter ratio in these mutants (51). In addition, many of the nucleotide synthesis pathway enzymes (PurACDF, GuaB, Prs, NrdA), including deoxynucleotide-producing RNR, have been localized to the inner membrane in a recent study (50). The replication protein interactants related to cell envelope metabolism identified in this study (Fig. 9B) may contribute to the already described regulatory mechanisms or represent alternative connections between cell membranes biology and DNA replication. In this work we confirmed that the DNA replication profile is perturbed in a mutant strain encoding one of the SeqA interactants – RfaD (Fig. 8), localized to the cytoplasmic side of the inner membrane (50). The mechanism of rfaD impact on chromosomal DNA replication control and the reason for appearance of asynchronous origin firing remain to be established. It is also worth noting that one of the interactants of DiaA protein found in this study, GalU, is an enzyme responsible for synthesis of UDP-glucose (Fig. 3, Supplementary file 2). This nucleotide sugar is necessary for production of cell envelope components including the LPS core. Intriguingly, that metabolite has also been implicated in regulation of the major division protein FtsZ, by binding to a moonlighting enzyme OpgH, involved in synthesis of osmoregulated periplasmic glucans (40). This way, the level of UDP-glucose influences *E. coli* cell size. Considering our results, it is possible that UDP-glucose metabolism is also implicated in DNA replication control.

**Figure 9.**
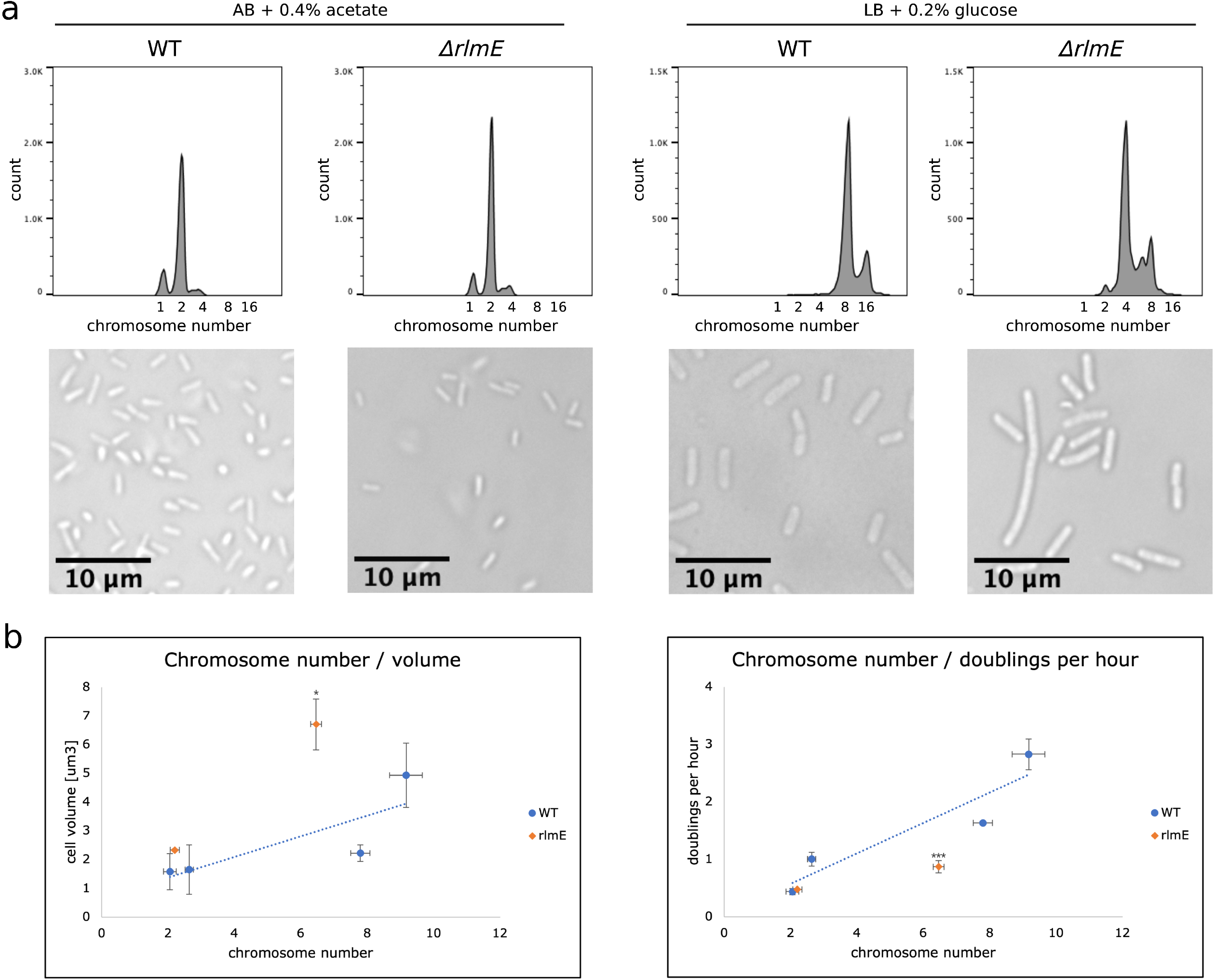
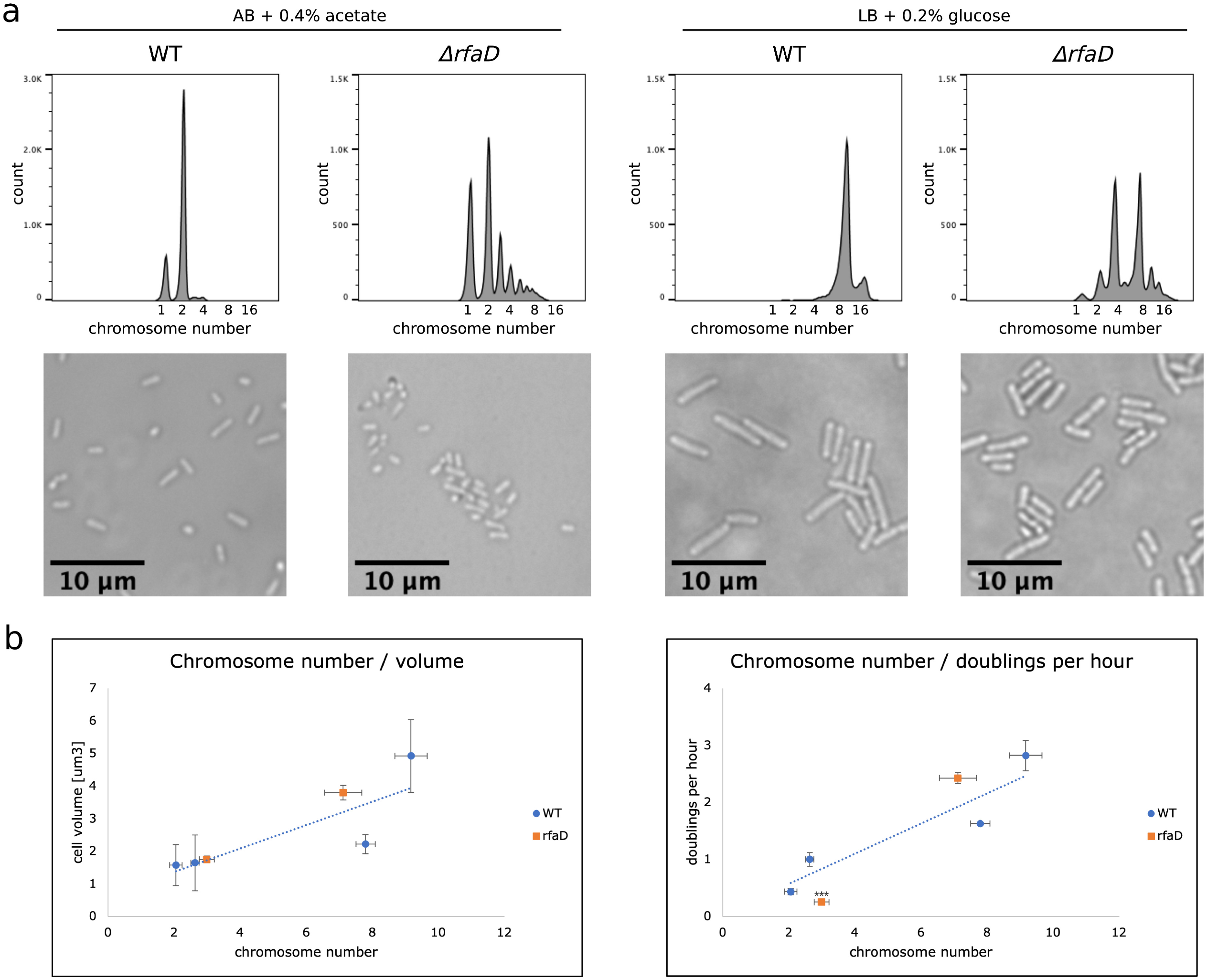
Functional connections between cell envelope biogenesis and DNA replication identified so far in E. coli (A). Cell envelope structure and genes engaged in biogenesis of particular layers (B). Proteins identified in this work as interactants of the replication proteins were highlighted in green.

Several reports have also suggested more or less direct regulation of various DNA replication-related protein activities by metabolic enzymes of the central carbon metabolism (CCM)(52– 57). Here, we observed several metabolic enzymes among replication protein interactants, with FbaB-NrdB (all conditions) and PfkB-DnaA (growth on M9+acetate) seeming highly specific (Fig. 3, Supplementary file 2).

Further studies are needed to verify physiological role of the uncovered PPIs. Moreover, implementation of more sensitive methods is required to detect very weak interactions and accurately quantify PPI network dynamics. However, this study proves that protein complexes formed by the *E. coli* replication proteins undergo large reorganization under different conditions which (Fig. 3; Fig 5), very likely, largely contributes to coordination of DNA replication with other cellular processes.

## Funding

National Science Center (Poland) UMO-2014/13/B/NZ2/01139 (M.G.) and UMO-2016/23/N/NZ2/02378 (J.M-O)

## Supplementary figure legends

**Figure S1.**
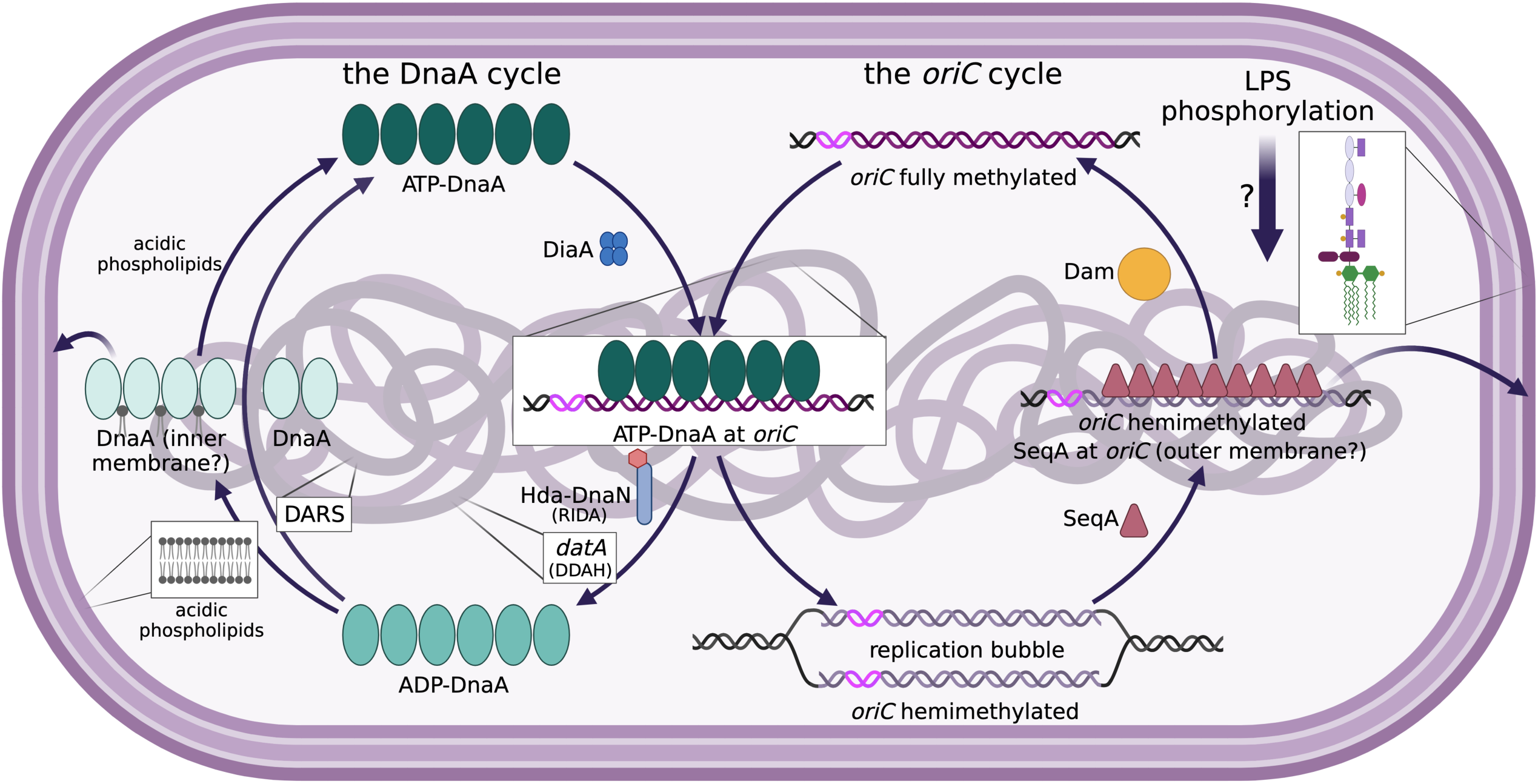
A-H. Similarity of three biological repetitions performed under each growth conditions for the analyzed bait proteins.

**Figure S2.**
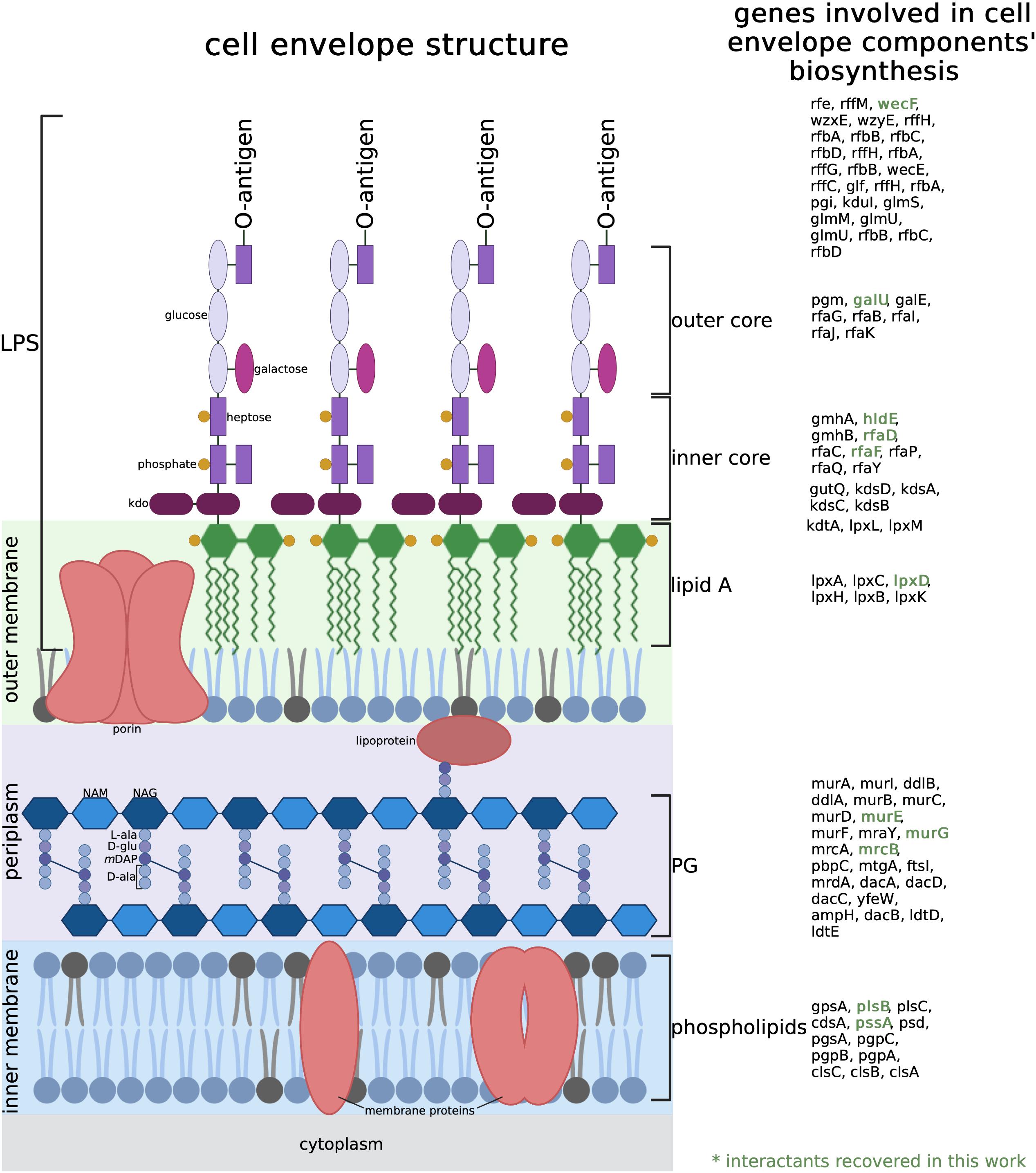
Flow cytometry results and standard curves depicting relation of chromosome number with cell volume and growth rate for the WT *E. coli* strain.

